# Integrative epigenetics and transcriptomics identify aging genes in human blood

**DOI:** 10.1101/2024.05.30.596713

**Authors:** Mahdi Moqri, Jesse Poganik, Chiara Herzog, Kejun Ying, Qingwen Chen, Mehrnoosh Emamifar, Alexander Tyshkovskiy, Alec Eames, Jure Mur, Benyamin Matei-Dediu, Ludger Goeminne, Wayne Mitchell, Daniel Mccartney, Riccardo Marioni, Jessica A. Lasky-Su, Michael P. Snyder, Vadim N. Gladyshev

## Abstract

Recent epigenome-wide studies have identified a large number of genomic regions that consistently exhibit changes in their methylation status with aging across diverse populations, but the functional consequences of these changes are largely unknown. On the other hand, transcriptomic changes are more easily interpreted than epigenetic alterations, but previously identified age-related gene expression changes have shown limited replicability across populations. Here, we develop an approach that leverages high-resolution multi-omic data for an integrative analysis of epigenetic and transcriptomic age-related changes and identify genomic regions associated with both epigenetic and transcriptomic age-dependent changes in blood. Our results show that these “multi-omic aging genes” in blood are enriched for adaptive immune functions, replicate more robustly across diverse populations and are more strongly associated with aging-related outcomes compared to the genes identified using epigenetic or transcriptomic data alone. These multi-omic aging genes may serve as targets for epigenetic editing to facilitate cellular rejuvenation.

## Introduction

Aging is associated with a myriad of molecular and cellular changes. Understanding the impact of these changes may provide insights into aging biology and help identify new therapeutic strategies to target aging and age-related disease. DNA methylation (DNAm) alterations rank among the most well-studied and prominent age-related molecular changes; importantly, these alterations generally replicate well across many different cohorts and species ^1^. DNAm is part of the epigenome and is involved in regulating gene expression, but the downstream consequences of most age-related DNAm changes remain incompletely understood, rendering it challenging to interpret their functional impact. Conversely, transcriptomic changes are more functionally informative than DNAm due to their indication of the current level of gene expression in tissues and cells. Age-related changes to the transcriptome have also been described and have facilitated the development of ‘transcriptomic clocks’ ^2^; however, these clocks generally feature lower replicability across cohorts and datasets than DNAm-based clocks ^3^, higher noise, and susceptibility to batch effects ^4^. These features make comparison of existing transcriptomic clocks across studies difficult.

The integration of multiple omics-based measures could assist in overcoming specific limitations of an individual readout. Previous studies have leveraged multi-omics analyses to derive DNAm-based predictors (or proxies) of features in other omics or clinical data, combining the reliability of DNAm profiling with the more functional insights derived from other molecular changes. For instance, DNAm-based “EpiScores” have been shown to predict plasma proteomic features ^5^, and recently a DNAm-based multi-omic aging clock was described ^6^. The combination of epigenetic and transcriptomic data has been suggested to improve biological age predictions ^2^, yet few studies to date have investigated direct links between age-related epigenetic and transcriptomic changes ^1,7–11^. These studies have reported conflicting findings, with some reporting that methylome changes are indicative of alterations in gene expression ^7–9^ and others noting limited association with expression of affected genes ^1,10,11^.

A comprehensive, integrative analysis of epigenetic and transcriptomic changes across the human lifespan has not yet been conducted. Such a study could shed further light on the functional relevance of age-related DNAm changes and identify genes that could be targeted to combat aging-related declines in physiological health. Here, we perform a large-scale analysis of DNAm and gene expression data from blood samples, utilizing data from several cohorts (n=4,174 and 3,461 total samples for DNAm and RNA-seq, respectively; **Table 1**). Our integrative multi-omic approach enabled the identification of functionally relevant DNAm changes in blood associated with gene expression alterations, hereafter referred to as “multi-omic aging genes”. We validate our findings in a new high-quality blood DNAm dataset, generated using the latest methylation array technology, from the Mass General Brigham Biobank (n=500) that comprises a broad range of ages and has additional data on many aging-associated outcomes. Our findings provide a first in-depth assessment of the functional consequences of age-related DNAm changes in blood, and highlight the potential for multi-omics approaches to uncover novel functionally relevant genes and genetic loci. In turn, these features could be used to develop meaningful predictors for relevant aging-related outcomes, and may be targeted by therapeutics to mitigate aging-related molecular changes and decline.

**Table 1.**
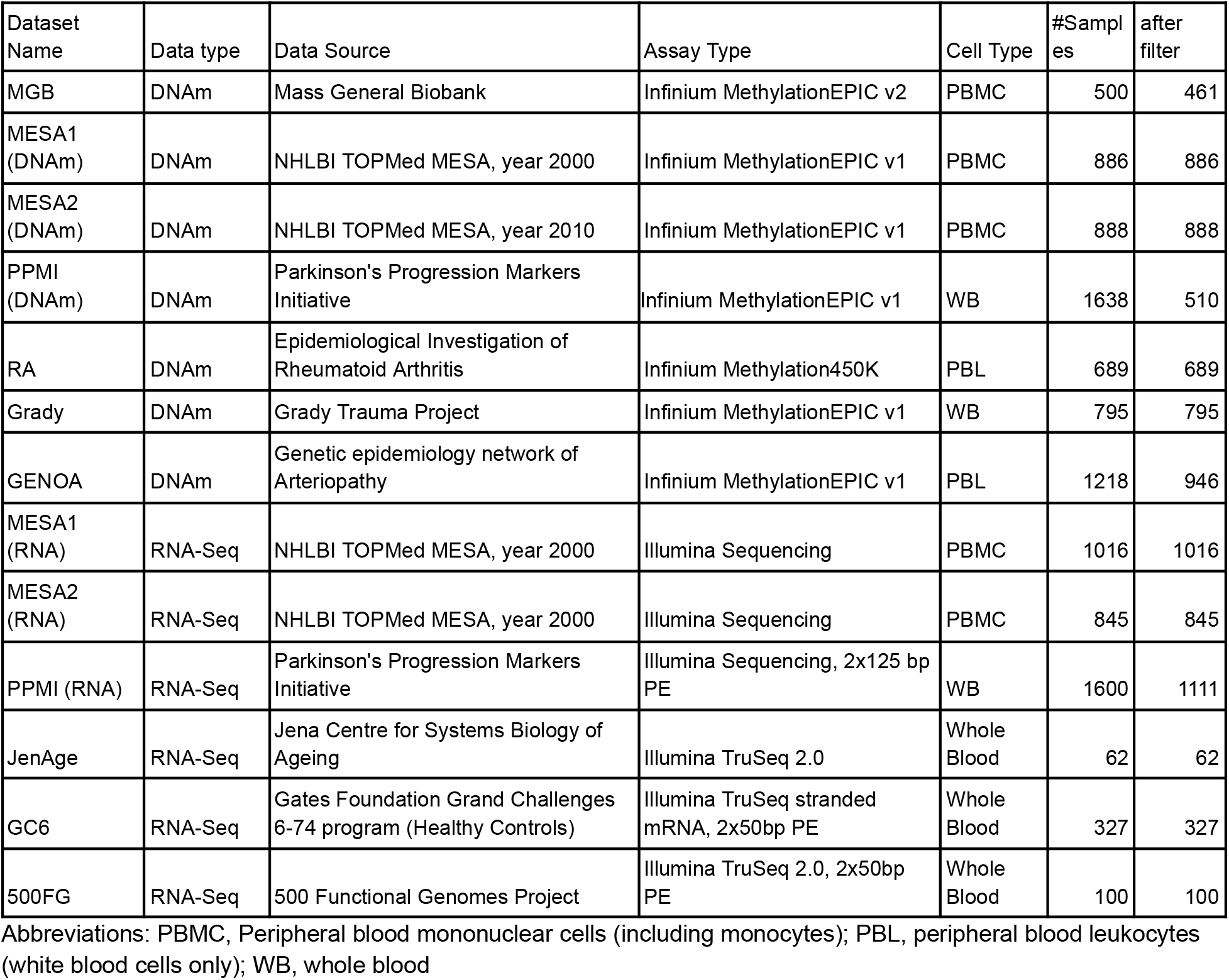
Datasets used in this study.

## Results

### Age-associated gene expression levels across studies and time points are less reproducible than DNAm patterns

We initially aimed to identify age-associated genes from six large transcriptomic datasets, including data from the Multi-Ethnic Study of Atherosclerosis (MESA), which features two sampling time points (referred to hereafter as MESA1 and MESA2); the Parkinson’s Progression Markers Initiative (PPMI); the Gates Grand Challenge (GC6); the 500 Functional Genomics Project (500FG); and the JenAge Ageing Factor Database. These datasets cover wide age ranges and are balanced between male and female participants (Fig. 1a). For each cohort, we assessed the correlation of gene expression levels with age (see Methods). Among the top 10 transcripts associated with age across all datasets, we observed that the correlation was quite variable between cohorts (Fig. 1b). For instance, we found loss of expression of CD248 as a function of age, consistent with previous reports ^12,13^, but the magnitude of the correlation ranged from -0.3 to -0.5 (Fig. 1c). For further validation of this observation, we examined the correlation between age and expression levels of “aging transcripts” previously identified by Peters et al. across these cohorts. In agreement with our previous result, we found generally low levels of correlation between expression levels of these genes and age across cohorts (Fig. 1d). Thus, age-related transcriptomic data appears to replicate poorly across cohorts.

**Figure 1.**
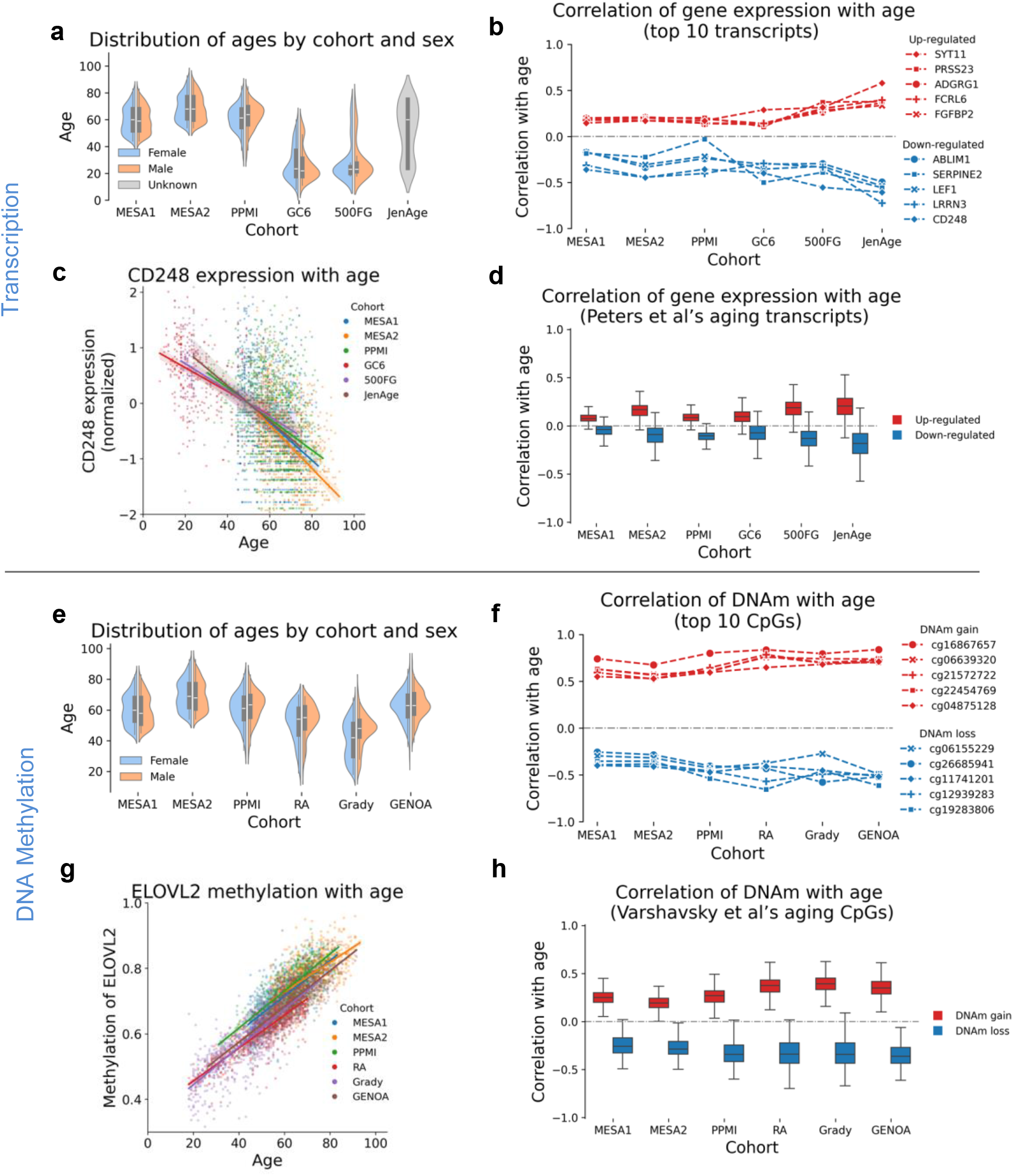
“Aging transcripts” and “aging CpGs” in blood. **a**. Distribution of ages by cohort and sex in 6 RNA-seq datasets used in this analysis. **b**. Top age-associated transcriptions across the cohorts. **c**. Gene expression levels of CD248 by age across the cohorts. d. Correlation between age and gene expression at “aging transcripts” (Peters et al 2015) across the cohorts. **e**. Distribution of ages by cohort and sex in 6 DNAm datasets used in this analysis. **f**. Top age-associated DNAm across the cohorts. h. DNAm levels of ELOVL2 promoter (cg16867657) by age across the cohorts. **g**. Correlation between age and DNAm levels at “aging CpGs” (Varshavsky et al 2023) across the cohorts.

We next asked whether the same was true of DNAm data. We analyzed DNAm from six large data sources: MESA1, MESA2, and PPMI, as above; a cohort of patients with Rheumatoid Arthritis and healthy controls (RA); the Grady Trauma Project (Grady); and the Genetic Epidemiology Network of Arteriopathy (GENOA) study (Fig. 1e). For the top 10 CpG sites that were found to be associated with age, we found considerably higher and more consistent correlations with age across the datasets (Fig. 1f). For instance, cg16867657, a well-known aging-associated CpG site within the *ELOVL2* promoter ^14^, emerged as the top site consistently positively correlated with age (Fig. 1g). Across “aging CpGs” reported by Varshavsky et al. in 2023 ^15^, we again found correlations that were larger in magnitude than those observed for aging transcripts, and more consistent across cohorts. Thus, age-associated DNAm data are considerably more replicable across cohorts than transcriptomic data.

### Integration of multiple omics modalities allows for the identification of validated age-associated genes

To identify age-associated genes that are consistent across blood samples of several cohorts, we integrated DNAm and transcriptomic data from studies that produced both data modalities (MESA and PPMI) to identify genes whose age-associated DNAm changes have functional consequences at the level of gene expression. On average, across MESA time points and across cohorts, we observed surprisingly little correlation between gene expression levels and age for genes whose CpG sites had the greatest gain of methylation over aging (Fig. 2a), consistent with some previous reports ^1,10^. Conversely, we found that genes with the greatest decrease in expression levels with aging consistently featured significantly higher DNAm correlation with age (Fig. 2b). These observations were consistent at the level of individual genes: There were minimal changes in the expression of *ELOVL2, KLF14*, and *FHL2*, the genes associated with the top-ranked aging-associated CpGs, between young and old individuals (Fig. 2c, *left*, see Methods for details). In contrast, the top down-regulated aging transcripts, CD248, LRRN3, and NELL2 each featured increased DNAm in older compared to younger individuals (Fig. 2c, *right*). CD428 in particular showed strong positive correlation with age at the level of DNAm (Fig. 2d, *left*) and strong inverse correlation with gene expression (Fig. 2d, *right*); this was true at both MESA time points and in the PPMI dataset. We were able to validate 106 such genes in all datasets (Fig. 2e). These genes feature age-dependent changes in both DNAm and expression levels; we term such genes *multi-omic aging genes*.

**Figure 2.**
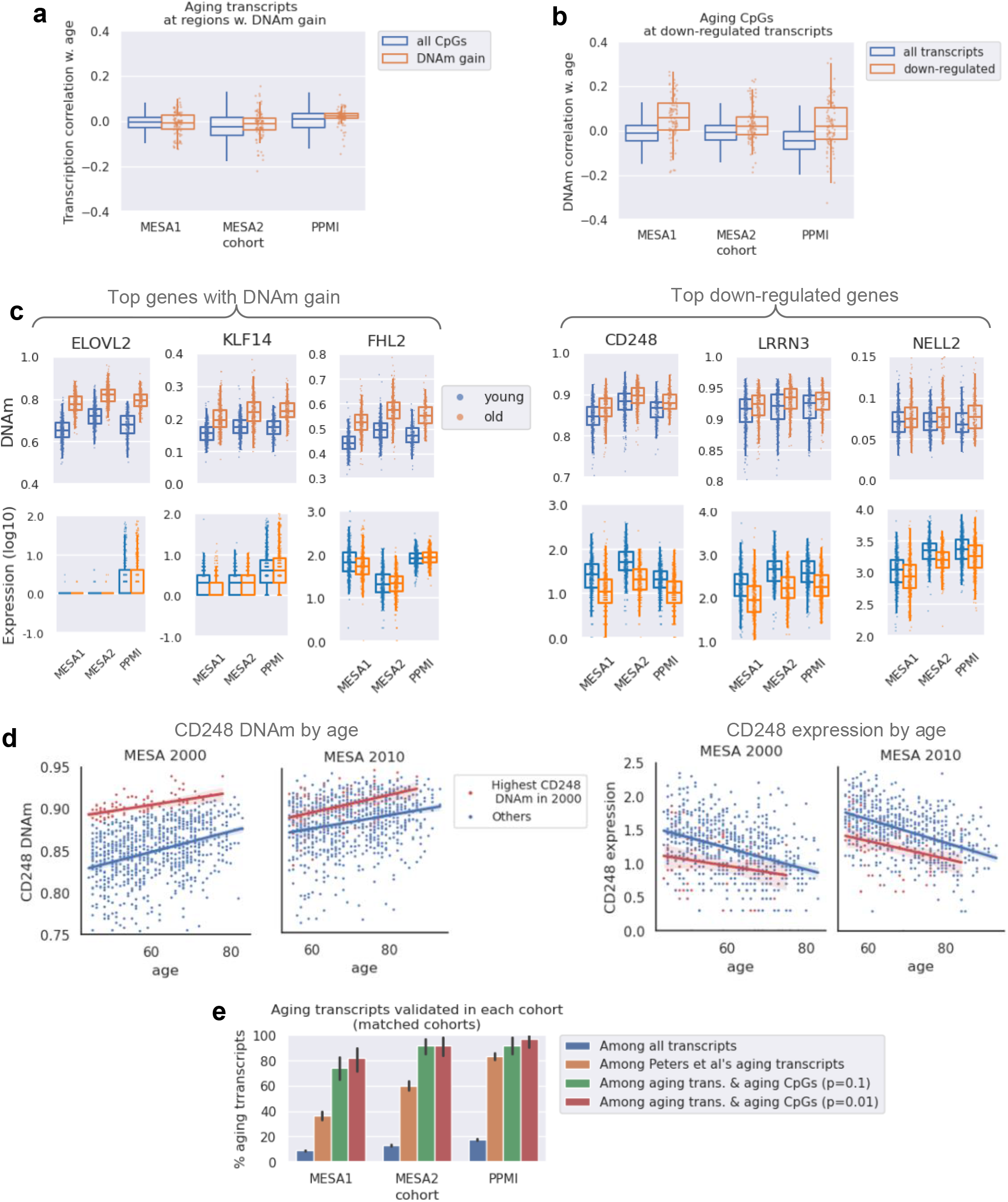
Multi-omics biomarkers of aging identify functional aging genes. **a**. Correlation of gene expression levels with age at regions with the highest age-associated DNA gain in MESA1, MESA2 and PPMI cohorts **b**. Correlation of DNAm levels with age at promoter of most age-associated down-regulated genes in MESA1, MESA2, and PPMI cohorts **c**. DNAm and expression levels of genes with the highest age-associated DNAm (left) and age-associated expression (right) levels for young (<=45 years) and old (>=75 years) participants in MESA1, MESA2, and PPMI cohorts **d**. DNAm and expression levels of CD248 in MESA1, MESA1 and MESA2 for individuals with top 10% highest DNAm (age-adjusted) at promoter of CD248 in MESA1 compared to others **e**. Percentage of top “aging transcripts” (Peters et al 2015) that replicates in MESA1, MESA2 and PPMI cohorts compared to the percentage of aging transcripts with corresponding aging CpGs that replicate in each cohort.

### Construction of a gold-standard reference DNAm dataset for aging human blood

To validate our novel multi-omic aging genes, we aimed to perform an independent external validation using a novel cohort. To do so, we generated DNAm profiles for 500 individuals of diverse ages from the Mass General Brigham (MGB) Biobank using the Illumina Infinium MethylationEPIC v2.0 array, which covers over 935,000 CpG sites enriched for regulatory regions and has been shown to exhibit high reproducibility ^16^. Our subjects were recruited from a major metropolitan academic medical center, were roughly balanced between male and female, and were generally representative of the racial/ethnic distribution of the local area (Fig. 3a).

**Figure 3.**
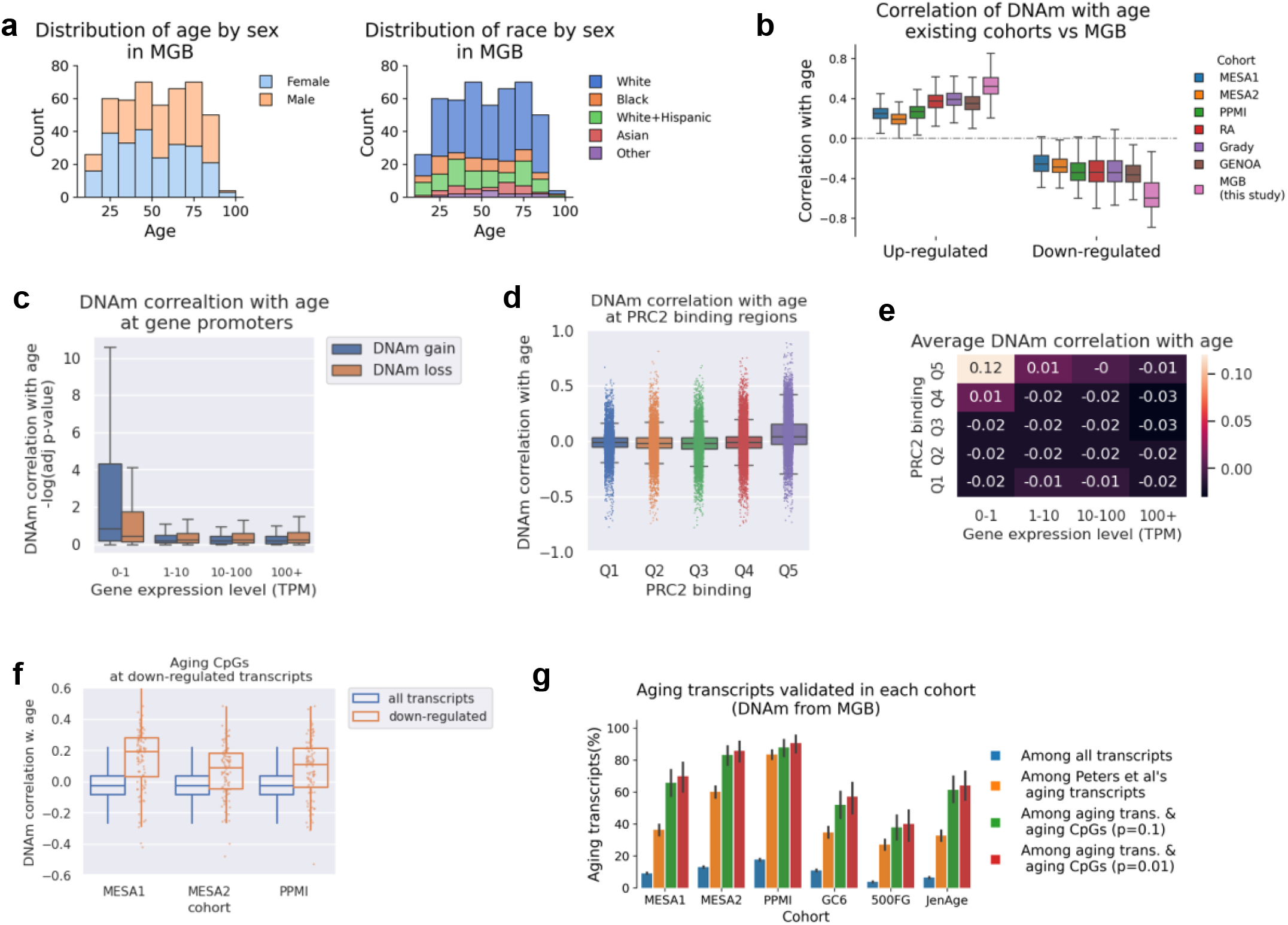
A reference epigenetic aging improves validation of transcriptomic aging. **a**. Distribution of sex (left) and race (right) by age in MGB cohort. **b**. Correlation between age DNAm levels at “aging CpGs” (Varshavsky et al 2023) in MGB cohort compared to other cohorts. **c**. Correlation between age and DNAm levels at promoters of genes with different levels of expressions. **d**. Correlation between age and DNAm levels at promoters of genes with different levels of PRC2 binding (Q5=highest 20%). **e**. Correlation between age and DNAm levels of genes with different gene expression and PRC2 binding levels. **f**. Correlation of DNAm levels with age (in MGB) at promoter of most age-associated down-regulated genes in MESA1, MESA2, and PPMI cohorts. **g**. Percentage of “aging transcripts” (Peters et al 2015) that replicates in each cohort compared to the percentage of aging transcripts with corresponding aging CpGs (based on MGB cohort).

As expected, DNAm levels at “aging CpGs” ^15^ exhibited consistent age-dependent changes, with slightly stronger correlations for the MGB cohort compared to other cohorts (Fig. 3b). Sites with the strongest DNAm correlation with age, particularly those that gain methylation, were enriched in lowly-expressed and repressed genes (Fig. 3c). Such genes were enriched for strong PRC2 binding (Fig. 3d, e). We also confirmed the gain of methylation at CpG sites within promoters of genes whose expression is inversely associated with age in this dataset (Fig. 3f). Finally, we again compared the concordance of the top age-associated genes identified by Peters et al. with gene expression data, with the addition of DNAm data from the MGB cohort. The addition of DNAm data considerably increased the percentage of age-associated genes we were able to validate in both MESA time points and in PPMI, as well as in the other three cohorts with RNA-seq data (Fig. 3g). Interestingly, we found our multi-omic aging genes to be particularly enriched for T cell-specific genes (e.g. CD27, CD28, CD248, TCF7). These analyses demonstrate that integration of multiple omics modalities has the ability to refine the identification and validation of true aging-associated genes. Moreover, our newly generated high quality dataset exhibits strong age-related DNAm changes, covers a broad age range, and features a multitude of aging-associated outcome data such as mortality. Thus, this dataset is ideal for benchmarking existing and novel biomarkers of aging.

### Multi-omic aging genes are predictive of aging outcomes

To evaluate the functional relevance of our newly identified multi-omic aging genes, we analyzed the extent to which they are associated with aging outcomes, particularly mortality. We utilized large datasets containing both DNAm data and mortality data: a larger subset of subjects from the MGB Biobank (MGB-4K), and Generation Scotland (GS). CpG sites associated with our multi-omic aging genes were strongly associated with mortality risk in both cohorts (Fig. 4a), with hazard ratios for the top CpGs ranging from approximately 1.3–1.75 per standard deviation (Fig. 4b). Importantly, survival analysis based on the top mortality-risk-associated CpG site showed clear stratification of subjects (Fig. 4c). Additionally, disease gene network over-representation analysis of our multi-omic aging genes revealed enrichment for numerous aging-associated diseases, particularly those related to the aging immune system such as various lymphomas (Fig. 5c). Thus, multi-omic aging genes are strongly associated with clinically-relevant aging-related outcomes such as disease and mortality risk.

**Figure 4.**
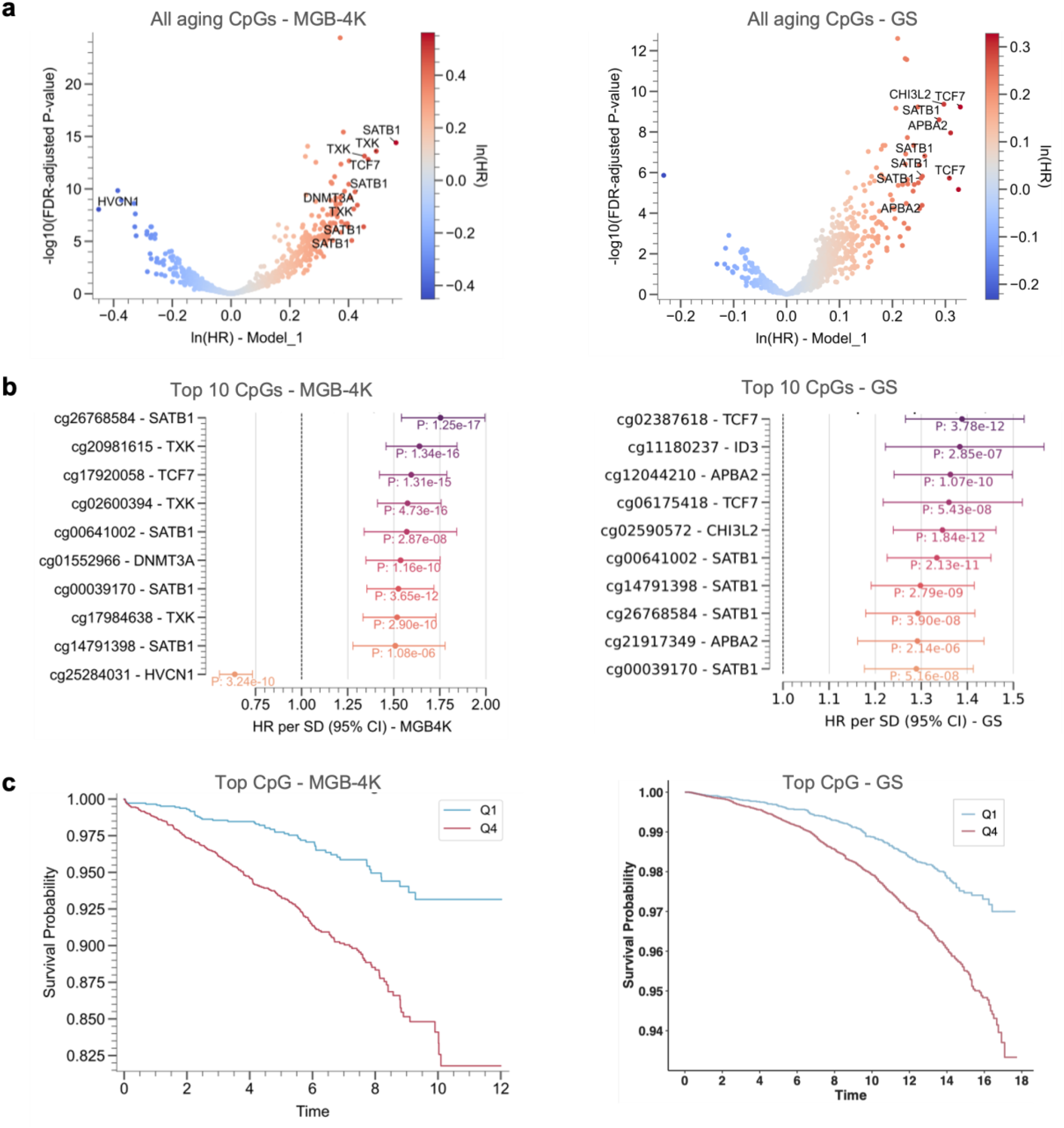
Association of “multi-omic aging genes” with mortality. (a) Volcano plot showing DNA methylation level predicting mortality in the MGB4K and GS cohorts, with top 10 CpGs labeled. (b)Hazard ratios (HR) per 1 standard deviation with 95% confidence intervals for the top 10 CpGs associated with mortality in the two cohorts. (c) Kaplan-Meier curves showing the association between the top CpG in each cohort and all-cause mortality.

**Figure 5.**
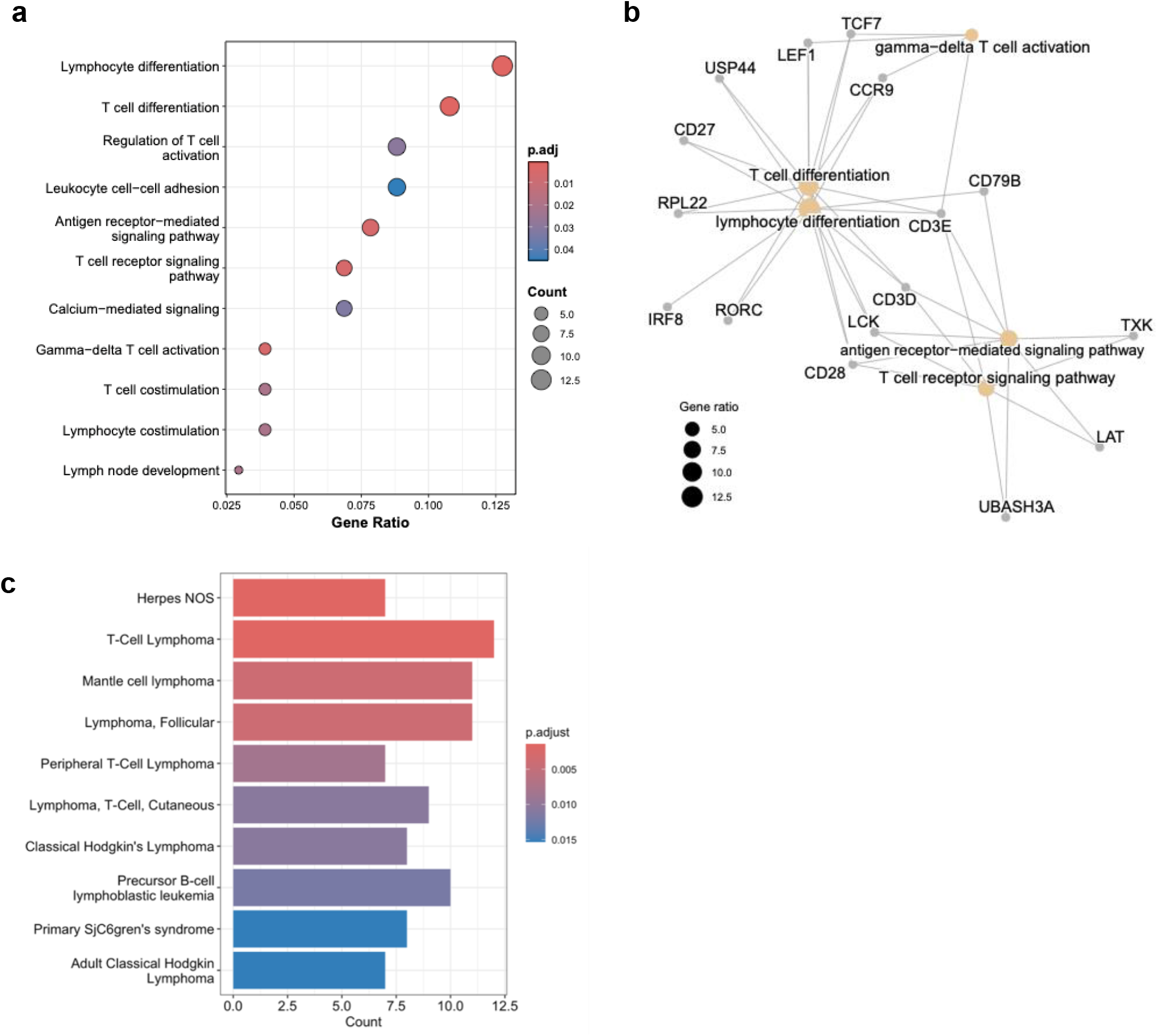
Association of aging genes methylation with cell composition. **a**. Functional enrichment of genes associated with aging at the level of gene expression and DNA methylation in blood. Size of dots reflects the number of genes, while color denotes statistical significance (BH-adjusted p-value). Fisher exact test based on GO BP ontology was used for enrichment analysis. Redundant functions with high semantic similarity were filtered out with the simplify method from clusterProfiler package. **b**. Gene-concept network of multi-omic blood biomarkers of aging. Gray dots correspond to genes, while yellow dots correspond to enriched functions. Genes associated with the corresponding terms are connected by edge. Size of the functional dot reflects the gene ratio of the corresponding term. **c**. Over-representation analysis for the disease-gene associations network.

### Age-related methylation changes at CpGs within multi-omic aging genes are largely independent of cell composition changes

Blood is a heterogeneous tissue composed of multiple cell types, the proportions of which change with age. Given these age-dependent alterations in blood cell composition, age-related changes observed in the transcriptome and epigenome may potentially be related to changes in cell proportions over time ^17^. Noting a clear enrichment for T cell differentiation genes in our multi-omic aging genes (Fig. 5a, b), we sought to investigate whether aging patterns of our multi-omic aging genes were truly intrinsic to individual blood cell types or simply a reflection of age-related cell composition changes. To do so, we generated methylation-based cell composition estimates ^18,19^ for our MGB cohort samples (Fig. 3a) using predictors of proportions for the most abundant blood cell types and ordinal abundances values for less common, yet age-dependent, cell types. We observed modest age-related changes in cell composition for the most common cell types (Figure removed—see Acknowledgement). However, for naive T cells, we found a dramatic decline associated with age that is consistent with a well-documented feature of human immune aging ^20^. Finally, we characterized the relationship between age and methylation level for CpGs within our identified multi-omic aging genes after adjusting for cell composition. Significant associations of methylation levels at these CpGs with age remained after the adjustment (Figure removed—see Acknowledgement), indicating that age-related changes in these CpGs are independent of changes in cell composition. Thus, methylation levels at CpG sites within our multi-omic aging genes are robustly associated with aging.

## Discussion

As omic profiling becomes increasingly common in cohort studies of aging, integration across multiple datasets and omics modalities has the potential to reveal molecular changes associated with aging at a level of resolution previously inaccessible. In the current study, we provide one of the most comprehensive integrative multi-omic analyses of age-related DNAm and gene expression changes to date, leveraging DNAm and RNA-seq data from over 4,600 and 3,500 samples, respectively, drawn from diverse cohorts. Additionally, we performed a fully external validation of our findings in a new, high-quality reference DNAm dataset for aging biomarkers (Figure 3) derived from a representative cohort in the Mass General Brigham Biobank. We expect this high-quality DNAm dataset will prove to be a useful future reference dataset to benchmark DNAm-based biomarkers of aging.

Our analysis revealed that CpGs most strongly correlated with aging do not necessarily predict changes in the expression of their associated genes. For instance, the *ELOVL2* gene that harbors the CpG most strongly correlated with age (Figure 1g) and included in the models of many population biomarkers of aging does not exhibit gene expression changes with age (Figure 2c). This observation applies to most of the top age-related DNAm loci (Figure 2a, c). This suggests that age-dependent gain of DNAm may occur at sites where cells can afford such changes because they do not lead to any functional consequences. Further work will be needed to identify the features that characterize such sites. Importantly, these results indicate that previous bioinformatic analyses of genes harboring such CpG sites should be interpreted with caution, as downstream ramifications of these DNAm changes may be limited.

To overcome this issue, we integrated DNAm and RNA-seq data to identify functionally relevant “multi-omic” aging genes in blood. Because this analysis explicitly coupled age-related changes in DNAm to changes in gene expression, the results are more biologically interpretable than those derived from DNAm data alone. Aging genes identified using our multi-omic approach are enriched for genes associated with lymphocyte and T cell differentiation and activation (e.g., *TCF7, CD27, CCR9, IRF8*, among others; Figure 5a), in line with the notion that age-dependent changes in T cell biology represent a prominent feature of aging ^21^. Interestingly, it has previously been suggested that T cell aging could play a role in whole-body deterioration, indicating that our newly identified genes may represent important targets for strategies to combat age-related disease. While aging also results in well-described changes in immune cell proportion ^22^ (Figure 5c), our analysis suggests that for the majority of our multi-omic aging genes, the association with age remains even after correction for cell type proportions (Figure 5d). Our findings also indicate that in contrast to age-related genes identified via RNA-seq alone, multi-omic aging genes much more robustly replicate across datasets (Figure 2e) and thus have the potential to improve reliability of transcriptomic clocks, which to-date have suffered from poor replicability across datasets.

While the correlation with aging was interesting, we were particularly motivated to explore the ability of our multi-omic aging genes to predict aging outcomes. A mortality risk analysis in several independent cohorts demonstrated that CpG methylation levels at sites associated with our multi-omic aging genes are highly predictive of aging outcomes (Figure 4a, c), with hazard ratios of individual sites of up to 1.58 per standard deviation (e.g., cg06175418 in *TCF7* Figure 4b). Thus, these results suggest that integration of multiple omics allows for the identification of reliable and interpretable aging genes and CpGs that associate with aging outcomes.

Most importantly, this study demonstrates that integrative multi-omic analyses have the powerful ability to overcome limitations associated with analyses of single omic modalities. In this sense, multi-omic analysis represents an important new frontier for the aging biology field. We anticipate that our findings will catalyze future integrative analysis and allow both for the development of improved and functionally interpretable biomarkers of aging using DNAm or transcriptomic data, and the identification of putative genetic targets for future interventions.

## Methods

### Study datasets

The current study leverages several previously-described DNA methylation and RNA sequencing datasets, shown in **Table 1**. Their underlying cohorts or population characteristics have been described in detail in publications by the original respective authors. Additionally, we generate a new DNA methylation dataset encompassing 500 individuals of various ages derived from the Mass General Brigham Biobank. This study was approved by the Mass General Brigham IRB (protocol 2021P003059). Details on methylation data and RNA sequencing data processing are described below. For comparison of molecular data between young and old groups, participants 45 years old or younger are labeled as “young” and those 75 years or older are labeled as “old.”

### Methylation data generation and processing

Previously processed described datasets were obtained from relevant sources (e.g., GEO, dbGaP, or others) and no further processing or filtering was conducted. New methylation data from the MGB biobank were generated using the Illumina HumanMethylationEPIC version 2 array (cat# 20087709), encompassing over 950.000 CpG sites. Raw .IDAT files were preprocessed using standard parameters in the R SeSAMe package ^23^, version 1.22.1.

### RNA seq data and processing

We obtained previous RNA-seq datasets as raw gene expression counts. Count data were normalized using log transformation on (raw counts + 1) and median adjusted.

### Survival analysis

In a previously described dataset (MGB-4k, referred to in Chen et al. 2023 ^6^), we applied a multivariate Cox Proportional Hazard regression model to test the association between all-cause mortality and each “aging-CpG” site, adjusting for age and sex. Next, we ranked the “aging-CpG” sites by Benjamini-Hochberg-corrected p-values from smallest to largest and selected the top 10 sites, as demonstrated in the volcano plot. For these top “aging-CpGs,” we drew a forest plot to show the point estimates and 95% confidence intervals of the adjusted hazard ratios for all-cause mortality. Additionally, we plotted the adjusted survival curves for these top 10 “aging-CpGs.”

### Cell composition analysis

To estimate the cell composition of each sample in the MGB cohort, we applied software provided by the Clock Foundation that deconvolutes cell composition using bulk methylation profiles. This software estimates the proportion of cell types through constrained quadratic programming for common blood cell types (CD8 T cells, CD4 T cells, NK cells, B cells, monocytes, and granulocytes) ^19^ and penalized regression for rarer cell types (plasmablasts, CD8+CD28-CD45RA-T cells, naive CD8 T cells, and naive CD4 T cells) ^18^.

For each CpG site associated with a “multi-omic aging gene”, we corrected for cell composition through the following approach: first, we applied multivariate linear regression to predict the methylation beta value of that CpG site using, as inputs, age and the cell proportion of each cell type, excluding granulocytes and plasmablasts due to high collinearity (defined as a variance inflation factor over 5). Next, we determined whether the regression coefficient p-value for age was still significant (Benjamini-Hochberg-corrected p-value < 0.05) in predicting CpG methylation.

### Functional enrichment analysis

Functional enrichment of genes associated with aging across modalities was performed using Fisher’s exact test and gene terms from GO Biological Process ontology with clusterProfiler package in R. Redundant functions with high semantic similarity were filtered out with simplify function. Adjustment for multiple comparisons was performed with the default Benjamini-Hochberg approach, and gene terms with adjusted p-value < 0.05 were considered statistically significant. Enriched functions were visualized with dot plot. Gene-concept network of multi-omic blood biomarkers of aging and representative enriched functional terms was constructed with the “cnetplot” function from “enrichplot” package in R. Over-representation analysis for the disease gene network (Fig. 5c) was conducted using R version 4.3.3 and the enrichDGN function from the DOSE package. This function leverages DisGeNET (Janet et al., 2015) to construct disease-associated gene networks.

## Acknowledgements

We thank the staff of the Mass General Brigham Biobank for assistance with sample procurement. Funded by grants from NIA and the Hevolution Foundation. Some figures using data from MGB500 are removed due to active competition on epigenetic biomarkers of aging hosted by the Biomarkers of Aging Consortium: https://www.agingconsortium.org/challenge

## References

1. Lu, A. T. et al. Universal DNA methylation age across mammalian tissues. Nature Aging 3, 1144–1166 (2023).

2. Peters, M. J. et al. The transcriptional landscape of age in human peripheral blood. Nature Communications 6, 8570 (2015).

3. Jylhävä, J., Pedersen, N. L. & Hägg, S. Biological Age Predictors. EBioMedicine 21, 29–36 (2017).

4. Meyer, D. H. & Schumacher, B. BiT age: A transcriptome-based aging clock near the theoretical limit of accuracy. Aging Cell 20, e13320 (2021).

5. Gadd, D. A. et al. Epigenetic scores for the circulating proteome as tools for disease prediction. eLife 11, e71802 (2022).

6. Chen, Q. et al. OMICmAge: An integrative multi-omics approach to quantify biological age with electronic medical records. bioRxiv 2023.10.16.562114 (2023) doi:10.1101/2023.10.16.562114.

7. Hannum, G. et al. Genome-wide methylation profiles reveal quantitative views of human aging rates. Mol Cell 49, 359–367 (2013).

8. Horvath, S. et al. DNA methylation aging and transcriptomic studies in horses. Nature Communications 13, 40 (2022).

9. Reynolds, L. M. et al. Age-related variations in the methylome associated with gene expression in human monocytes and T cells. Nature Communications 5, 5366 (2014).

10. Dozmorov, M. G. Polycomb repressive complex 2 epigenomic signature defines age-associated hypermethylation and gene expression changes. Epigenetics 10, 484–495 (2015).

11. Steegenga, W. T. et al. Genome-wide age-related changes in DNA methylation and gene expression in human PBMCs. AGE 36, 9648 (2014).

12. Nakamura, S. et al. Identification of blood biomarkers of aging by transcript profiling of whole blood. Biochemical and Biophysical Research Communications 418, 313–318 (2012).

13. Calabria, E. et al. Aging: a portrait from gene expression profile in blood cells. Aging (Albany NY) 8, 1802–1817 (2016).

14. Garagnani, P. et al. Methylation of ELOVL2 gene as a new epigenetic marker of age. Aging Cell 11, 1132–1134 (2012).

15. Varshavsky, M. et al. Accurate age prediction from blood using a small set of DNA methylation sites and a cohort-based machine learning algorithm. Cell Reports Methods 3, 100567 (2023).

16. Noguera-Castells, A., García-Prieto, C. A., Álvarez-Errico, D. & Esteller, M. Validation of the new EPIC DNA methylation microarray (900K EPIC v2) for high-throughput profiling of the human DNA methylome. Epigenetics 18, 2185742 (2023).

17. Jaffe, A. E. & Irizarry, R. A. Accounting for cellular heterogeneity is critical in epigenome-wide association studies. Genome Biology 15, R31 (2014).

18. Horvath, S. DNA methylation age of human tissues and cell types. Genome Biol 14, R115 (2013).

19. Houseman, E. A. et al. DNA methylation arrays as surrogate measures of cell mixture distribution. BMC Bioinformatics 13, 86 (2012).

20. Lazuardi, L. et al. Age-related loss of naïve T cells and dysregulation of T-cell/B-cell interactions in human lymph nodes. Immunology 114, 37–43 (2005).

21. Mittelbrunn, M. & Kroemer, G. Hallmarks of T cell aging. Nature Immunology 22, 687–698 (2021).

22. Nikolich-Žugich, J. The twilight of immunity: emerging concepts in aging of the immune system. Nature Immunology 19, 10–19 (2018).

23. Zhou, W., Triche, T. J., Laird, P. W. & Shen, H. SeSAMe: reducing artifactual detection of DNA methylation by Infinium BeadChips in genomic deletions. Nucleic Acids Research (2018) doi:10.1093/nar/gky691.

